# Digital Phage Biology in Droplets

**DOI:** 10.1101/2025.01.14.632946

**Authors:** Louis Givelet, Sophie von Schönberg, Florian Katzmeier, Friedrich C. Simmel

## Abstract

Since their discovery, bacteriophages – viruses that infect bacteria – have been an invaluable source of insight for molecular biology and have provided a plethora of molecular components for biotechnology. Driven by the imminent global antibiotic crisis, interest in using bacteriophages as antimicrobial agents has strongly increased in the past few years. While standard quantification of phage counts and infectivity by double-layer plaque assays (DLA) have provided foundational insights, they are limited by their inability to monitor infection dynamics over time and the inflexibility in experimental setups. Here, we introduce a novel high-throughput method using droplet microfluidics to quantify individual phage infection events, enhancing the understanding of phage-host dynamics at the single-event level. By co-encapsulating individual phages and bacteria in microfluidic droplets, we can control key experimental parameters such as exposure time and the ratio of phages to bacteria. Our system allows for precise quantification of lysis events and the direct observation of lysis kinetics unaffected by the continuous release of progeny phages that is typical in bulk cultures. Moreover, our method is potentially applicable to any phage-host pair. Our findings suggest that droplet microfluidics can provide a more accurate and dynamic understanding of phage biology. This approach could be particularly valuable in the development of phage-based antimicrobial strategies in the face of rising antibiotic resistance.

---

Since the first discoveries of bacteriophages more than a century ago [1, 2], their continual study had a major impact on biological research, both at the fundamental level and in terms of applications. Being recognized as the most abundant biological entity on earth [3], phages are ubiquitously present in extremely diverse environments [4], typically outnumbering their bacterial hosts by an order of magnitude. Phages are pervasive to prokaryotic life and therefore exert significant selective pressure on microbial communities. Through co-evolution with their hosts, they increase the rate of molecular evolution in bacteria and play a pivotal role in horizontal gene transfer, thus substantially shaping the complexity of ecosystems [5–7].

Bacteriophages, as obligate parasites, exist in a metabolically inactive virion form outside of their host cells, entirely dependent on infecting bacterial hosts for replication. This fundamental simplicity has been instrumental in elucidating several key concepts in biology, and thus had a major impact in establishing the foundational principles of molecular biology [8]. Bacteriophage research has resulted in a multitude of applications in molecular biology and biotechnology, such as type II restriction enzymes [9], DNA ligases [10], phage display [11] or CRISPR-Cas tools [12].

Besides the immense biotechnological potential of phage biology, an emerging health crisis has renewed the interest in phage studies in the scientific community in recent years: the rise of antibiotic resistance has become a pressing public health concern, prompting warnings of the onset of a ‘post-antibiotic era’ [13–15]. Utilizing bacteriophages as therapeutic agents in combating bacterial infections is increasingly considered as a promising alternative to antibiotics [16–18]. Notably, this potential was already recognized shortly after the initial discovery of phages [2, 19], but could not be fully exploited so far.

One of the most robust methods to study phage-host interactions is the double layer plaque assay (DLA), used primarily for quantification of phages [20, 21]. In this technique, host cells and phages are incubated inside a thin layer of soft nutrient agar (usually atop a bottom layer of nutrient agar), allowing diffusion of phages as well as cell growth inside the agar layer. Deposited phages infect bacteria, replicate within them and are released after bacterial lysis. Repeated cycles of infection, replication and lysis result in the creation of plaques: circular lysis zones, visible as clear areas that are straightforward to count. The technique thus allows quantifying the virulence of a phage solution with respect to a specific bacterial host, measured in terms of ‘Plaque Forming Units’/mL (PFU/mL).

Even though DLA is considered the ‘gold standard’ for phage quantification since its invention in 1936 [20, 22], it suffers from a couple of significant drawbacks. DLA represents a fixed and rather inflexible experimental setup and can suffer from poor reproducibility [23, 24]. Successful titer determination is dependent on certain properties of both host and phage: the ability of the host to grow to sufficient density, as well as the release of an adequate number of progeny phages to form a discernible plaque. The dependence on host growth rate also dictates that - for the majority of bacterial hosts - DLA requires at least overnight incubation, for slow-growing strains it may even require incubation for several days [20, 24]. Additionally, the semi-solid agar environment not necessarily represents a physiological environment for all phage-host pairs. Traditional alternatives for the enumeration of phages are microscopy-based methods such as TEM or molecular methods like qPCR [20]. Lastly, DLA is not suited for kinetic studies of phage infection [24]. Indeed, the first time point where plaques can be identified is reached only when the bacterial host has grown to sufficient density and is close to stationary phase.

As a consequence, the dynamics of phage infection are usually not monitored over time using plaque assays, but rather in bulk liquid culture measuring optical density [25] or using fluorescent DNA dyes [24]. However, all of these methods characterize phage infection at the population level, where the continuous release of newly formed phages prevents the study of individual phage infection events.

In the present work, we develop a novel, statistics-guided approach towards the quantification of individual phage infection events using droplet microfluidics. Droplet microfluidics has emerged as an important enabling technique for a wide range of bioanalytical applications over the past decade [26–30]. Droplet microfluidics was employed to isolate and amplify phages [31–34], and various droplet-based phage detection and enumeration methods were implemented, relying on digital PCR [35], heterologous protein expression [36], DNA-intercalating dyes [33, 37], bacterial growth rate monitoring [38] or light scattering [39]. Except for recent work by Nikolic et al. [40], however, the potential for investigating bacteriophage infection dynamics in microfluidic droplets was not fully recognized yet.

While studying phage propagation and virulence in bulk environments can provide population-scale information, inferring the dynamics of individual phage-host interactions is extremely challenging. By contrast, droplet microfluidics is ideally suited to study thousands of individual phage-host interactions involving small numbers of phages and bacteria in parallel, independently and with minimal volume requirements. Our high-throughput approach allows to precisely control key parameters such as the time of exposure of bacteria to phages, the compartmentalization volume, and the ratio between phages and bacteria encapsulated. Furthermore, our setup is applicable to any phage-host pair and provides an absolute quantification of lysis events.

## Results

### A high-throughput digital phage assay

In all previous work that utilized droplet microfluidics for counting or analysis of bacteriophages, the phages were invariably mixed with their bacterial hosts in a single bulk suspension before they were compartmentalized into droplets for further study. While this approach is straightforward, it requires the rapid encapsulation of phages and bacteria after mixing, in particular when studying phages with a short lysis cycle such as T7 phage [41]. In our workflow, suspensions containing phages and bacteria are co-injected into a flow-focusing microfluidic chip to ensure that the exposure of the bacteria to the phages begins precisely at the moment of encapsulation (Figure 1a). For the detection of bacterial lysis events in the droplets, our method relies on the labeling of the bacterial DNA released after cell lysis by the cell wall-impermeable intercalating fluorescent DNA dye YOYO-1 (Figure 1b). Even though YOYO-1 is also capable of labeling phage DNA [33, 42], we expected droplets containing lysed hosts to display substantially higher fluorescence levels than droplets containing only phages due to the sheer size difference between the phage and bacterial genomes. We confirmed that this is indeed the case using bulk assays (see Figure S2). After collection and incubation of the droplet emulsions for a defined time interval, the emulsions are re-injected into a separate microfluidic screening chip and scanned at rates of up to several kHz using a dedicated high-speed fluorescence detection setup (Figure 1c,d). For each droplet, a red and green fluorescence emission signal is acquired. During production of the droplet emulsions, we introduce a red fluorescent reference dye (Atto 565), which allows to collect additional information on the droplets such as their size and the mixing fraction *α* between the phages and the bacteria. *α* is defined as the volume fraction of the phage suspension in the droplet, and thus the bacterial suspension has a mixing fraction of 1 *− α*. Based on the green fluorescence signal distribution caused by YOYO-1, droplets are digitized (defined as “positive” for lysed and “negative” for non-lysed or “1” and “0”, respectively) to determine the digital titer (Figure1e).

**Fig. 1.**
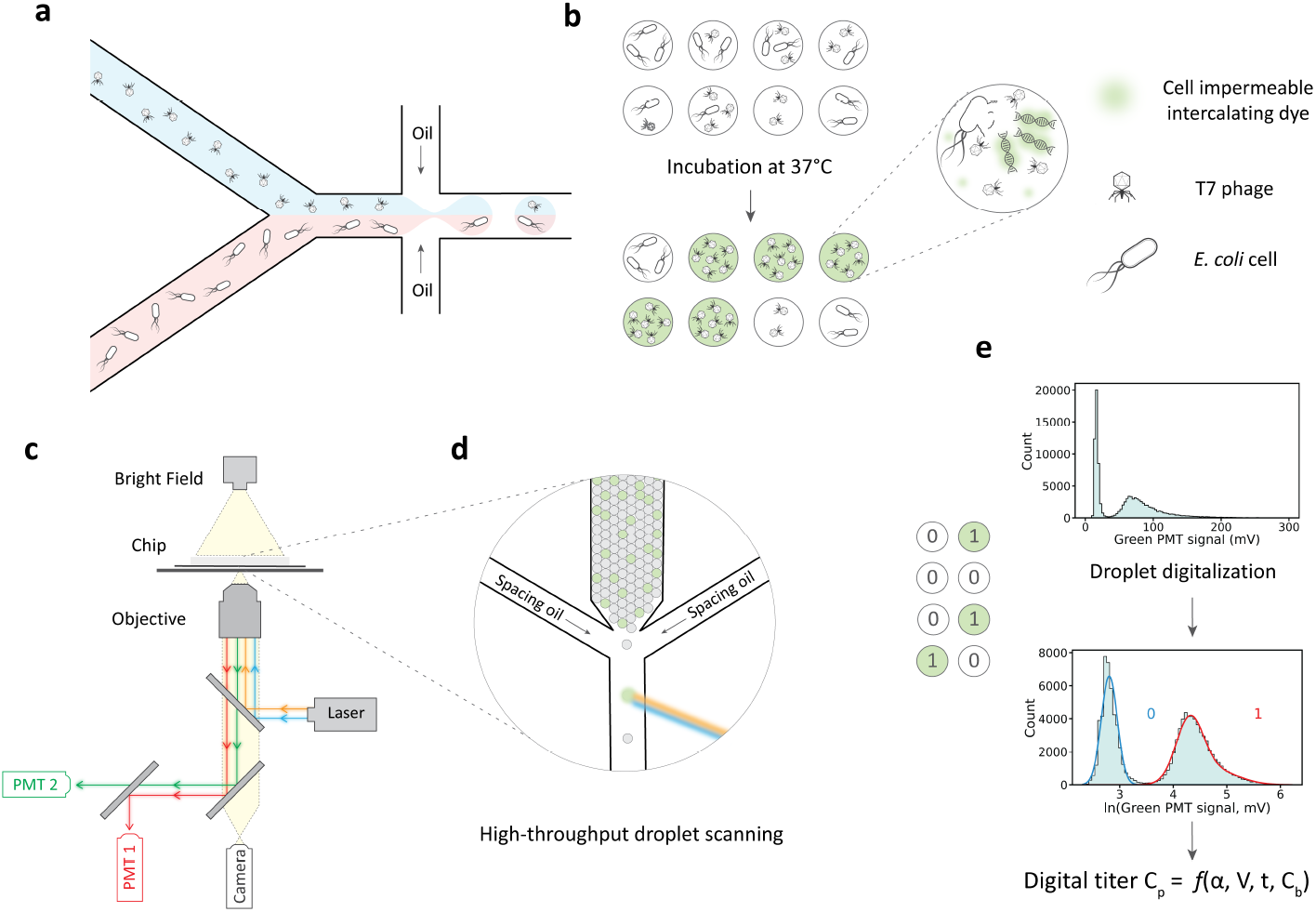
Experimental workflow.**(a)** Co-encapsulation of phages and bacteria, in presence of the DNA intercalating dye YOYO-1. **(b)** Reaction outcomes for different droplet compositions. **(c)** Experimental setup for droplet analysis. **(d)** Close-up schematic of the droplet analysis chip. The spacing oil ensures that the droplets are well separated and excited by the laser sequentially. **(e)** The green fluorescence signal (from YOYO-1) allows digitizing the droplets and computing the digital phage titer, taking into account various parameters such as the mixing fraction between phage and bacterial suspensions, droplet volume, time of incubation, and bacterial cell density.

### Statistics of bacterial lysis in droplets

To obtain the phage titer *c*_*p*_ of the encapsulated solution, we have to relate it to the fraction of positive droplets that exhibit a lysis event and thus become fluorescent. This depends on the statistics of the number of encapsulated phages and bacteria. To derive the probability *P*_*L*_ that a droplet exhibits a lysis event, we assume that every encapsulated phage will lead to the lysis of a bacterium when one is available. In other words, for a droplet to exhibit a lysis event, it must encapsulate at least one bacterium and at least one phage. Assuming independent encapsulation of each phage and bacterium, we can model the probability *P*_*b*_ and *P*_*p*_ of encapsulating exactly *b* bacteria and *p* phages within a droplet using Poisson distributions [43]. These distributions are each characterized by a single parameter, the expected numbers *λ*_*b*_ and *λ*_*p*_ of encapsulated bacteria and phages, respectively. *λ*_*p*_ can be calculated from the phage stock titer *c*_*p*_, the droplet volume *V*, and the mixing fraction *α* as *λ*_*p*_ = *αc*_*p*_*V*. Similarly, the expected number of encapsulated bacteria is given by *λ*_*b*_ = (1 *− α*)*c*_*b*_*V*, where *c*_*b*_ is the bacterial stock density. According to the Poisson distribution, the probability of having exactly zero phages encapsulated is *P*_*p*=0_ = exp(*−λ*_*p*_), and thus the probability of having at least one phage encapsulated is given by *P*_*p>*0_ = 1 − exp(*−λ*_*p*_). Similarly, the probability of having at least one bacterium encapsulated in the droplet is *P*_*b>*0_ = 1 − exp(*−λ*_*b*_). Thus, the probability *P*_*L*_ that a droplet contains at least one bacterium and at least one phage is given by

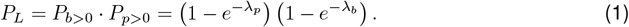

Based on our assumption, *P*_*L*_ corresponds to the fraction of droplets that are identified as positive, which is evidenced by their higher fluorescence signal. Equation 1 thus allows the calculation of the phage titer *c*_*p*_ for a given droplet volume *V*, bacteria density *c*_*b*_, and mixing fraction *α* from *P*_*L*_, which can be experimentally determined through the observation of a large enough number of droplets.

### Dynamic range of the digital titer

To test the functionality and performance of our system, we measured the green fluorescence levels of droplets of various compositions (Figure 2). When phages and bacteria were co-encapsulated, we observed a multimodal distribution for the green fluorescence signal, whereas distributions for droplets containing only bacteria (Figure 2e) or only reference dye (Figure 2f) were monomodal. The emulsion containing bacteria displays a slight extension towards higher fluorescence values, which is absent in droplets containing only reference dye. We attribute this minor increase to uncontrolled release of bacterial DNA, which is the result of unspecific cell death occurring during bacterial culture and preparation. During digitization, droplets belonging to the lower fluorescence mode are categorized as negative (‘0’), whereas droplets displaying a high level of fluorescence are classified as positive (‘1’).

**Fig. 2.**
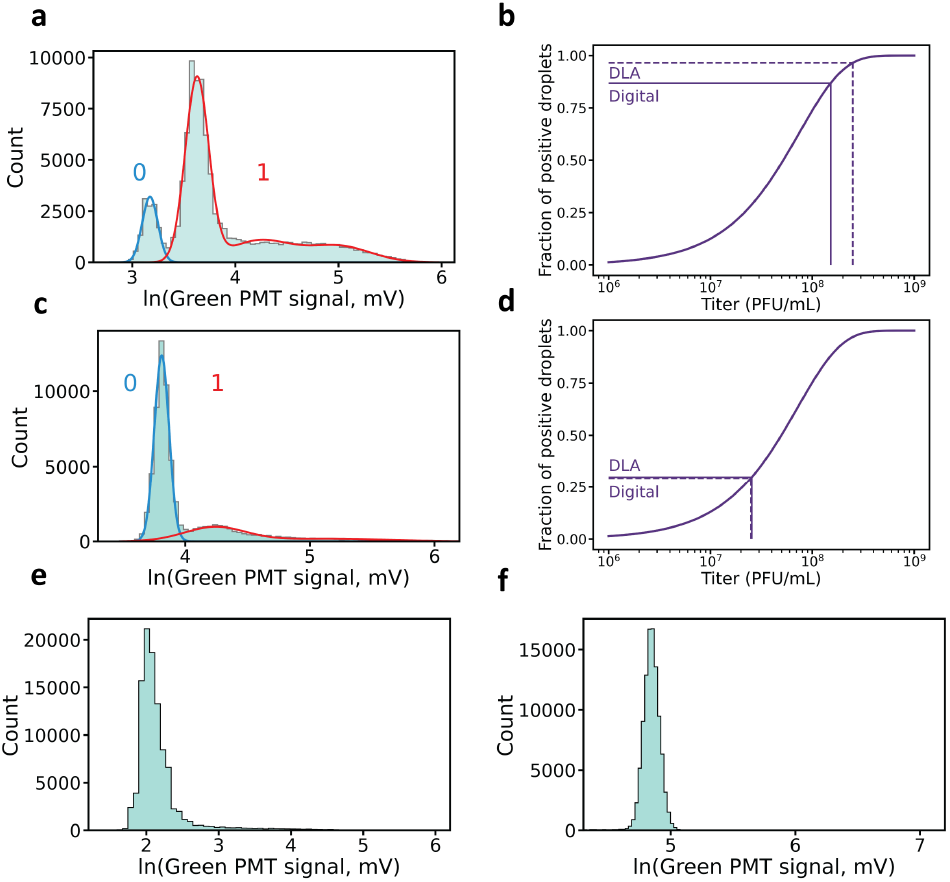
Droplet digitization and digital titer determination. **(a), (c)** Logarithm-transformed green fluorescence signal distribution (light blue bars). The modes detected by GMM are represented by blue (“0”, negative droplets) and red curves (“1”, positive droplets). *P*_*L*_ is determined by computing the sum of all of higher modes weights. **(b), (d)** relationship between phage titer and fraction of positive droplets *P*_*L*_. Vertical dotted lines indicate the titer determined by DLA. Horizontal dotted lines materialise the expected *P*_*L*_ based on DLA titer. Digital titers (vertical plain lines) are calculated from *P*_*L*_ (horizontal plain line). Their computed digital titers of 2.553 × 10^7^ and 1.516 × 10^8^ PFU/mL, lie close to the titers determined through DLA, 2.50 × 10^7^ and 2.50 × 10^8^ PFU/mL respectively. **(e)** logarithm-transformed green fluorescence signal distribution without phages. **(f)** logarithm-transformed green fluorescence signal distribution without bacteria nor phages.

Interestingly, encapsulating the suspension with the higher phage titer led to a bimodal distribution of the fluorescence signals for the positive droplets (Figure 2a). By contrast, droplets formed with the low-titer suspension resulted in a monomodal positive distribution (Figure 2c). This suggests that the higher positive fluorescence mode visible in Figure 2a represents droplets in which several lytic cycles took place, resulting in large amounts of free DNA in solution, a high phage count, and the lysis of multiple bacteria.

To achieve droplet digitization, we rely on a Gaussian Mixture Model (GMM), which enables automated separation of the droplet subpopulations. It also minimizes the requirement for user input and helps to avoid user bias in this critical step. The droplet fluorescence distributions can be well described by log-normal distributions, and we therefore apply the GMM to the logarithm-transformed green fluorescence signals. Through the GMM, one can determine the fraction of positive droplets *P*_*L*_, which then allows to determine the digital phage titer via Equation 1. The digital titers obtained in this way closely aligned with the titers determined by DLA.

As can be observed in Figure 2b & d, the relation between *P*_*L*_ and phage titer flattens out for very low and very large titers, suggesting an optimal intermediate range, where the two values are well correlated. Outside of this range, small changes in the fraction of positive droplets translate to large variations in the computed digital titer, making its determination unreliable. Notably, as our system allows the analysis of large numbers of droplets (10^4^-10^6^), even extremely unbalanced proportions between positive and negative droplets can be theoretically determined with strong statistical confidence. Nevertheless, digital titers lying outside this dynamic range will inevitably suffer from a higher imprecision. This is especially true for small titers, where unspecific bacterial lysis, as observed in Figure 2e, can lead to false positives.

### Screening with discrete mixing fractions

We investigated next whether the dynamic range of the digital titer could be expanded by analyzing droplets with different mixing fractions *α*. To this end, we dynamically changed the pressures at the two inlets of the aqueous solution while maintaining a constant overall dispersed phase pressure to ensure a uniform droplet size. Subpopulations of droplets with five different *α* values were generated and then identified via the fluorescence level of the red reference dye as described above (Figure 3a). The numerical values for *α* were determined by analyzing the corresponding microscope images (Figure 3b).

**Fig. 3.**
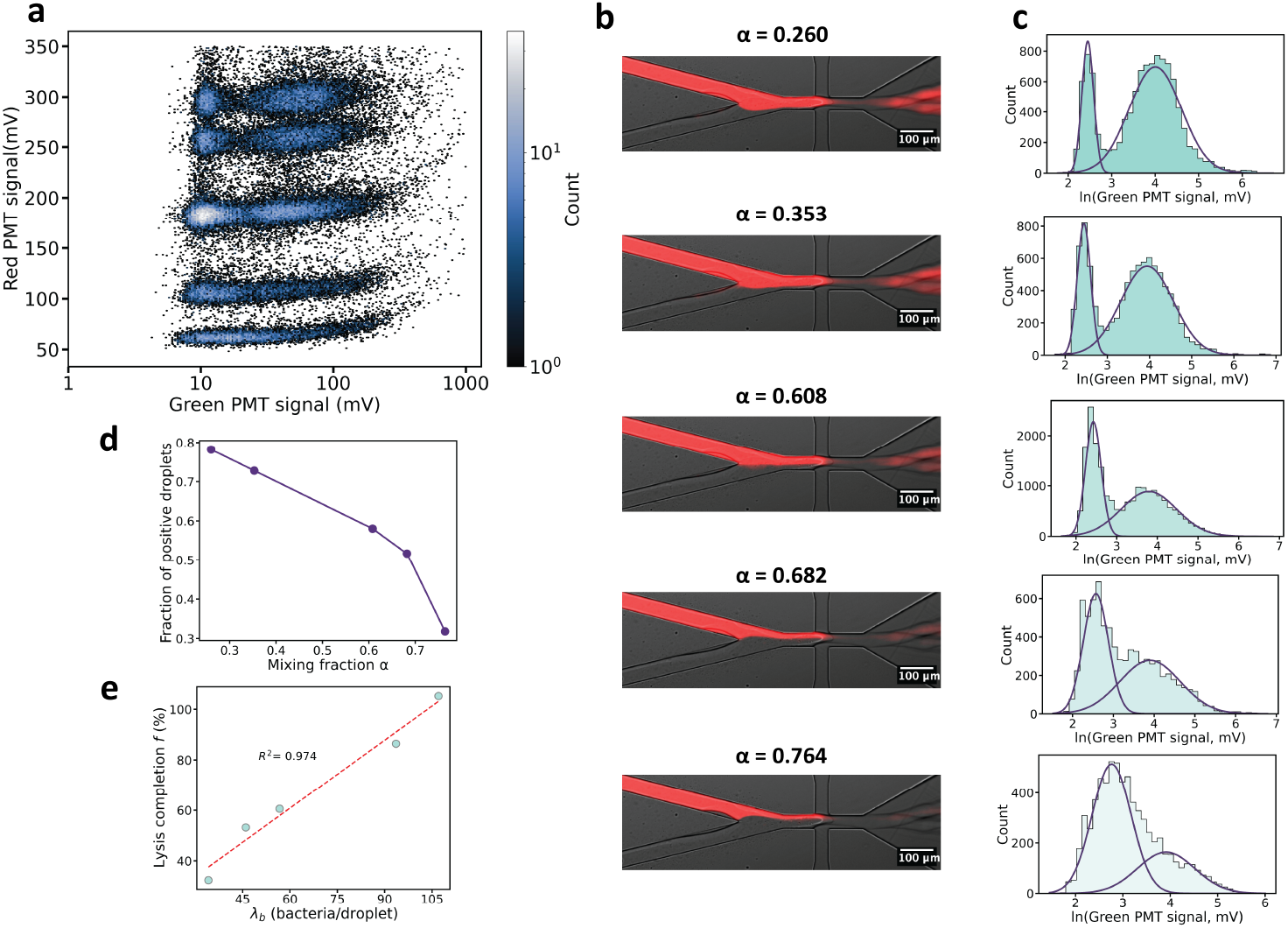
Screening of discrete mixing fractions. **(a)** 2D histogram plot of red and green fluorescence signals. **(b)** Fluorescence and bright-field microscopy overlay image of the flow focusing junction during droplet generation. The determined average value of the mixing fraction *α* is indicated for each subpopulation. **(c)** Logarithm-transformed green fluorescence signal distribution for each subpopulation (light blue bars). Purple curves represent the positive and negative droplet modes predicted by GMM with respective silhouette scores of 0.603, 0.625, 0.608, 0.630 and 0.620. **(d)** fraction of positive droplets *P*_*L*_ in function of *α*.**(e)** Lysis completion in function of the expected bacteria droplet occupancy *λ*_*b*_. The 100% lysis completion fraction of positive droplets is computed from DLA titer and *λ*_*b*_ from bacterial cell density in solution. dotted lines: linear fit.

A combined analysis of the subpopulations theoretically results in an expansion of the dynamic range. Smaller mixing fractions *α* are more appropriate for determining high titers, whereas larger mixing fractions *α* are better suited for estimating low titer values (see Figure S4a). As expected, the five *α*-subpopulations displayed distinct green fluorescence signal distributions and different fractions of positive droplets (Figure 3c). However, the quantitative influence of the mixing fraction *α* on the observed fraction of positive droplets was surprising. In our experiment, we intentionally used a high bacterial density, ensuring that the expected number of bacteria in the droplets for all tested mixing ratios satisfies *λ*_*b*_ *>* 30. This leads to a situation where virtually every droplet contains at least one bacterium for all mixing ratios *α*. Consequently, the fraction of positive droplets *P*_*L*_ is expected to depend solely on the expected number of phages encapsulated. As the expected number of phages per droplet, *λ*_*p*_, is directly proportional to *α*, droplets with a larger *α* were assumed to exhibit a higher fraction of positive droplets compared to those with a smaller *α*. Contrary to this expectation, the fraction of positive droplets was observed to decrease with increasing *α*, as shown in Figure 3d.

To explain this result, we hypothesize that the expected number *λ*_*b*_ of encapsulated bacteria might be lower than anticipated based on OD measurements of the initial bacterial suspension. This indicates that *λ*_*b*_ should be interpreted as the *effective* number of bacteria capable of participating in a lysis reaction, excluding those that are dead or in a dormant metabolic state [44, 45]. Furthermore, we suspect that the fraction of positive droplets also depends on the kinetics of the lysis reaction. Phage adsorption, and thereby lysis, occurs more slowly at lower bacterial densities [46], corresponding to higher mixing fractions *α*. Thus, at high fractions *α*, the lysis reaction may not yet have been completed.

We quantified the degree to which the observed fraction of positive droplets, 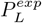, deviated from the theoretically expected fraction, 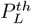, at the time of observation by computing the fraction of lysis completion as 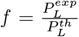. We computed 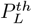 using equation 1, where we used a phage titer *c*_*p*_ measured via DLA, a cell density *c*_*b*_ derived from the optical density (OD) of the encapsulated bacteria solution, and the given values of the mixing fraction *α* and the droplet volume *V*. Notably, the parameter *f* (Figure 3e) appears to be (linearly) correlated with the expected number of bacteria per droplet, hinting at a critical influence of the bacterial count on the speed at which lysis occurs, or an overall lower effective cell density.

### Screening with continuous mixing fractions

We next sought to explore whether we could determine the effective bacterial density within the experiment and thereby achieve a more accurate assessment of the digital titer. In particular, by continuously varying *α* rather than in discrete steps, we can obtain a large dataset of different *P*_*L*_ values for various mixing fractions *α*, which allows for the confident determination of both bacterial density *c*_*b*_ and phage titer *c*_*p*_. To this end, the mixing fraction was constantly changed during the emulsification process, resulting in monodisperse droplets with a large range of *α* values (see Figure S5). To avoid the potential influence of lysis kinetics on the results, bacterial density was substantially increased compared to the previous experiments.

We binned droplets according to their red fluorescence value, and linearly mapped these bins to specific *α* values, using the minimum and maximum *α* measured through microscopy as reference points (see SI). For each bin, our GMM method was used to determine *P*_*L*_ which allowed us to determine its dependence on *α* (Figure 4). Fitting our statistical model (Eq. 1) for the fraction of positive droplets *P*_*L*_(*α*) to the experimental data yielded a digital titer (see Figure 4) of 1.205 × 10^8^ PFU/mL, which is very close to the titer (1.180×10^8^ PFU/mL) determined by DLA, thus validating our approach as a viable alternative to the traditional method. Notably, the effective bacterial density determined via the fit was found to correspond to only around 16.6% of that calculated from an OD measurement in bulk. As mentioned earlier, we hypothesize that the effective bacterial concentration excludes all bacteria that are not susceptible to infection for various reasons, including dormant metabolic states, resistance, and potentially dead cells or debris that contribute to the optical OD [44, 45].

**Fig. 4.**
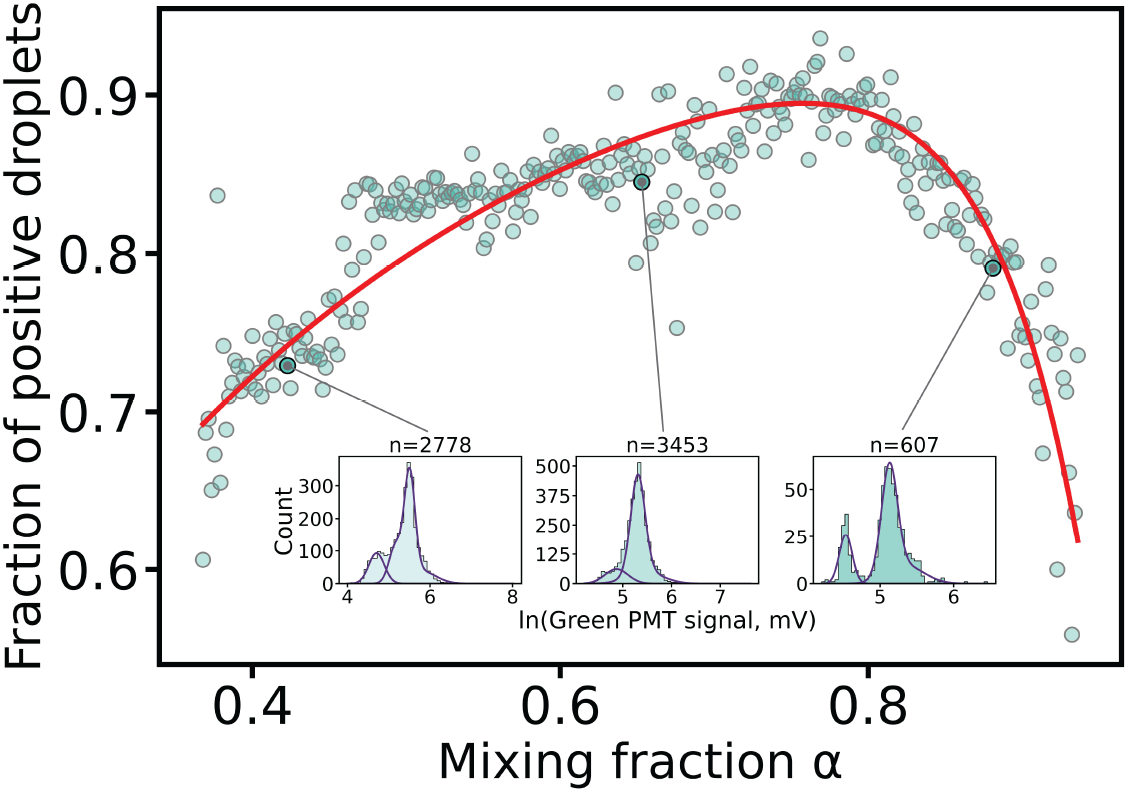
Continuous screening of the mixing fraction. Blue dots represent red fluorescence signal bins, containing each *≈* 2400 droplets on average. The total droplet population is 7.20 × 10^5^ droplets. Each bin is assigned to a *α* value according to its mean red fluorescence signal and its fraction of positive droplets *P*_*L*_ is determined by GMM. Insets used here as examples, represent the logarithm-transformed green fluorescence signal values distribution (blue bars) for three bins across the *α* values range as well as the modes determined by GMM (purple curves). The red curve represents a fit function of the equation 1 (*R*^2^ = 0.784) leading to a phage titer of 1.205 × 10^8^ PFU/mL and on OD of 0.773.

### Lysis kinetics in emulsions with a bimodal size distribution

Finally, we utilized our system to study the kinetics of phage infection by monitoring co-encapsulated phages and bacteria over time. To investigate how lysis kinetics depends on droplet size, we assessed the fraction of positive droplets for two different droplet volumes. By varying the pressure applied to the dispersed phase, we generated two distinct droplet populations within the same emulsion (Figure 5a). The resulting populations substantially differed in mean diameter, but each in itself was monodisperse (Figure 5b). The two populations and larger droplets resulting from droplet coalescence could be clearly separated from each other via the red reference fluorescence signal (Figure 5c). Similar to emulsions containing subpopulations with different mixing fractions *α*, the combined analysis of different droplet sizes enables an extension of the titer dynamic range (Figure S6).

**Fig. 5.**
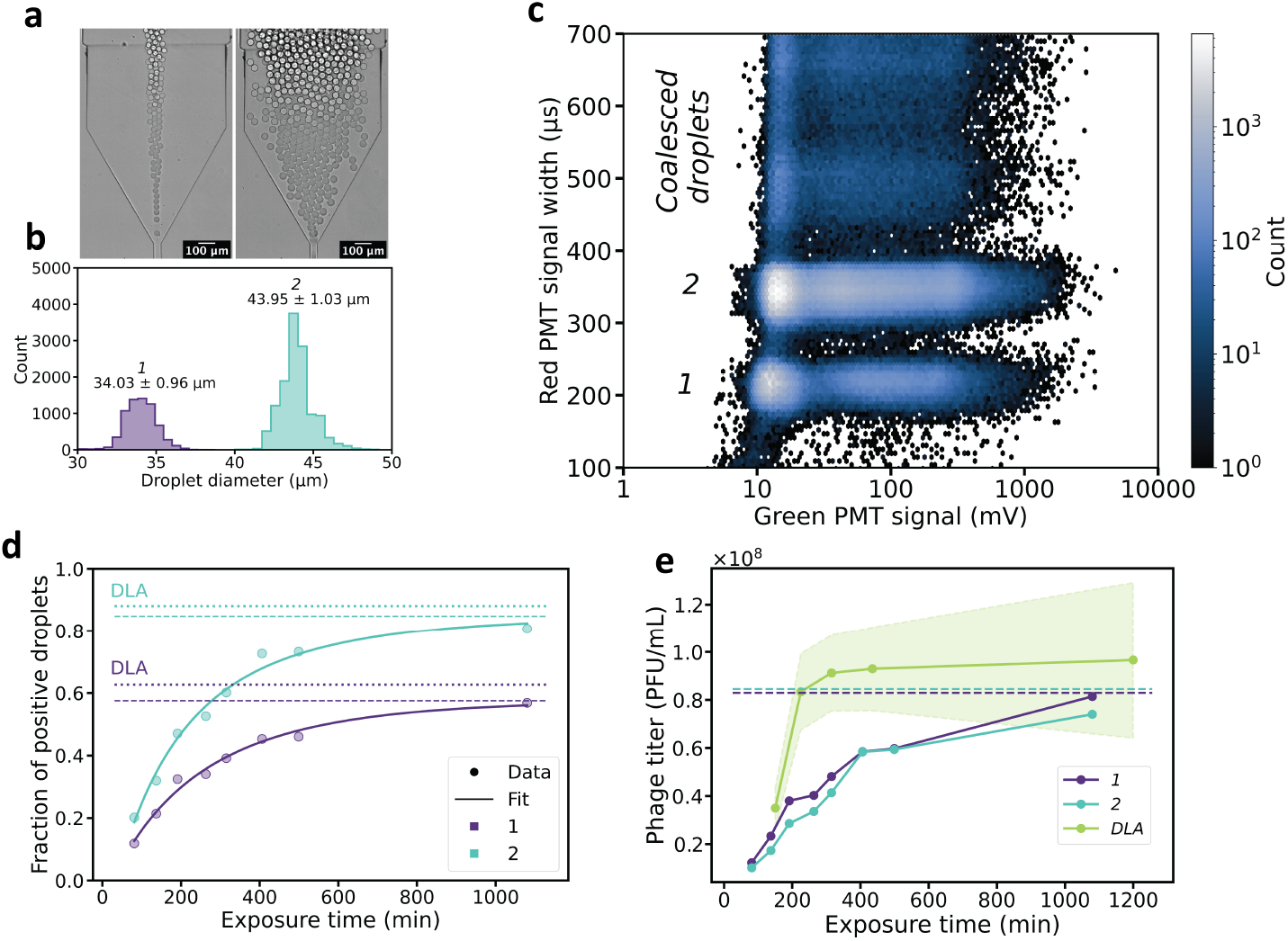
Analysis of emulsions with a bimodal size distribution. **(a)** Monitoring droplet generation by bright field microscopy. **(b)** Droplet size measurements. **(c)** 2D histogram plot of green fluorescence signals and red fluorescence signal width after 191 min of incubation. **(d)** fraction of positive droplets from different exposure time of bacteria to phages for each size mode. Dots: data from droplet digitization. Curves: fit function of Equation 2. Dashed line: limit value from fit. Dotted line: computed fraction of positive droplets based on titer obtained from DLA.**(f)** Digital phage titer computed from different exposure times for each size mode and phage titer from DLA. The shaded area represents the standard deviation from 3 technical replicates. Dashed line: phage titer from fit for 2 size modes.

The bimodal emulsions were analyzed after different incubation times, and for each subpopulation, the temporal evolution of the fraction of positive droplets was obtained (Figure 5d). Both droplet sizes exhibited similar lysis kinetics, asymptotically approaching a plateau value that nominally corresponds to the successful completion of lysis in all droplets containing both phages and bacteria. Supporting this interpretation, the fraction of positive droplets computed from the DLA titer after overnight incubation precisely aligns with this value.

To gain further insight, we developed a baseline model for phage infection kinetics in droplets (cf. Supplementary Information for details). In the model, phage adsorption follows second-order mass action kinetics, characterized by a rate constant *k*, consistent with the commonly employed bulk model [46–53]. A droplet is classified as positive in our model if at least one phage adsorbs to a bacterium. Furthermore, we incorporate a time delay *τ* between encapsulation and the detection of the first positive droplets, which can be interpreted as a maturation time [53, 54]. Assuming a relatively large number of encapsulated bacteria, in line with the experimental conditions, the fraction of positive droplets *P*_*L*_(*t*) at time *t* is given by

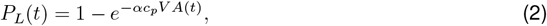

where

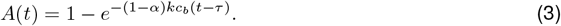

Here, the product *kc*_*b*_ can be interpreted as an effective first-order rate constant [46]. Fitting the equation to the data also allowed us to determine the phage titer, resulting in 8.30 × 10^7^ PFU/mL and 8.46 × 10^7^ PFU/mL for the smaller and larger droplets, respectively. The maturation times were *τ* = 24.6 min and *τ* = 34.5 min, and the effective first-order rate constants were *kc*_*b*_ = 0.0061 min^−1^ and *kc*_*b*_ = 0.0050 min^−1^. As before, the digital titers closely matched the DLA titer, which was determined to be 9.55×10^7^ PFU/mL. Furthermore, the determined lag times *τ* are of the same order of magnitude as previously reported values for T7 phage maturation [41]. However, the lag time *τ* is also affected by the time required for emulsification (which takes up to 10 minutes). The rate constant *k* for T7 phage adsorbing to *E. coli* ranges from 1 × 10^−10^ mL min^−1^ to 1 × 10^−9^ mL min^−1^ for different media [50]. With a bacterial density *c*_*b*_ of 3.52 × 10^9^ cells/mL, we determined the adsorption rate constant as *k* = 1.041 × 10^−11^ mL min^−1^ for the smaller droplets and 8.536 × 10^−12^ mL min^−1^ for the larger ones. These calculations took into account the effective bacterial fraction of 16.6% determined earlier. We attribute the significant deviation of the adsorption rate to the fact that we used PBS and not cell culture media in our experiment [50], and also to the imprecise determination of *c*_*b*_ in optical density measurements. The time courses depicted in Figure 5d suggest that droplet volume does not significantly affect the time scale of lysis. Notably, our model accurately accounts for the effect of varying droplet sizes, as evidenced by the similar fit parameters obtained for the data from the two volumes.

Remarkably, the time required to lyse all ‘lysis competent’ bacteria differs considerably between the digital assay and DLA (Figure 5e), for which lysis is completed much more rapidly. In a double-layer agar plate assay, complete lysis is defined as the point at which every phage deposited in the agar has resulted in the formation of a visible plaque. Lysis completion is not influenced by the progeny of the initial phage as the plaques are spatially isolated from each other. The faster transition to lysis completion in DLA can likely be attributed to the fact that each phage in the DLA is surrounded by bacteria in a growth supporting medium, which is critically influencing the phage infection dynamics. [20, 55]

## Discussion

In this work, we introduced a novel high-throughput technique based on droplet microfluidics that facilitates the study of phage infections in a digitized manner. Unlike other established methods, our approach is potentially applicable to the study of essentially any pair of a lytic phage and its host. Moreover, since the composition of the droplet can be selected freely, it enables tuning of experimental parameters such as the composition of media or microbial densities. Additionally, the temperature and duration of phage exposure can be freely adjusted. Droplet microfluidics allows the encapsulation of host-phage couples at different mixing fractions into hundreds of thousands of droplets within a very short period of time, enabling high-throughput analysis of individual phage infection events with high statistical confidence.

We demonstrated several variations of a droplet-based method that can provide comprehensive insights about the phage-host interactions through the digitization of infection events. The method enables the determination of phage titers with high accuracy. Modulating the mixing fraction of the phage and bacterial suspensions allows to adjust and expand the dynamic range of phage lysis experiments to a broad range of PFU values. Furthermore, this approach provides extensive information beyond that of single average composition emulsions. This can be exploited to determine the effective titers of both participants of the lysis reaction, host and phages.

Moreover, our system can be utilized to study phage infection kinetics at the single-event level. In conventional bulk assays, the study of infection kinetics is complicated by the fact that progeny phages from each burst are released to the environment and thus contribute to the infection of the remaining bacteria, masking the effect of the initial phage population. By contrast, encapsulation of phages and their hosts into small compartments allows counting discrete infection events, which can be attributed to the initially present phages. In other terms, infection kinetics in a bulk assay is dominated by the most infectious phages in the suspension and their progeny, while our digital assay gives insight into the diversity of phage infectivity within the population.

In the present study, we deliberately encapsulated bacteria in PBS, which does not support growth. This simplifies analysis as the experiments are performed with a constant bacterial count in each droplet. However, our system could also be used with culture media in which the bacteria grow to their maximum density, which would reduce the time required for the completion of lysis in all droplets [20, 55]. Our experiments could be performed with potentially up to 10^7^ droplets, which would allow the study of systems with extremely unbalanced ratios between positive and negative droplets. This would enable to identify very subtle differences of infectivity between phage-host pairs. Finally, our droplet microfluidic setup can be easily interfaced with a droplet sorting unit, which enables screening and selection of phage variants of interest. This capability could be highly beneficial for applications in phage engineering and directed evolution, offering a promising approach to optimize therapeutic phages for maximum infectivity and a broader host range. In conclusion, the digital phage experiments presented in this study enable the determination of multiple key infectivity parameters in a single experiment. Our method thus holds potential for widespread applications, ranging from basic phage biology research to the development of novel therapeutic phages.

## Methods

### Preparation of microfluidic devices

Microfluidic chip designs were developed using AutoCAD software. Silicon master molds were created from 2-inch silicon wafers (Siegert Wafer, Germany), employing photolithography with SU8-3050 photoresist (Micro resist Technology) and a μMLA100 tabletop maskless aligner (Heidelberg Instruments). Subsequently, PDMS microfluidic devices were crafted via soft lithography. Specifically, 15 g of PDMS (Sylgard 184, Dow Corning) was thoroughly mixed with 1.2 g of curing agent, poured over a silicon master enveloped in aluminum foil, degassed for 30 minutes, and then cured in an oven at 80°C for one hour. After curing, the PDMS was removed from the master, trimmed to size, and inlet and outlet holes were introduced using a biopsy punch. Glass slides were cleaned with 2% Hellmanex III (Hellma) and distilled water, followed by drying at 80°C for 30 minutes. Finally, the PDMS devices and glass slides were treated with O_2_ plasma (1 minute, 20 sccm, 100W) for bonding, then further cured together at 80°C for an hour.

### High-throughput production of droplets

Bacterial cell cultures were prepared via dilution (1/500) of a saturated pre-culture, incubated at 37°C, 250 rpm in NZYCM medium for 2 hours, then centrifuged (3000 G, 5 min) and re-suspended at an adequate bacterial density in PBS buffer (Fisher Scientific), typically *OD*_600_ > 2.5. One solution containing either phages or bacteria was supplemented with 2 μM of a reference dye (Atto 565, Sigma-Aldrich). Both solutions were supplemented with 2 μM of YOYO-1 (Invitrogen). For continuous mixing fraction *α* screening, the concentrations YOYO-1 and Att0565 were 1 and 4 μM, respectively. Microfluidic channel walls were treated with 1% trichloro(1H,1H,2H,2H-perfluorooctyl)silane (Merck) dissolved in FC-40 oil (Merck) for 1 min immediately prior to use. Microfluidic devices were operated using a 4-channel Elveflow OB1 controller for the aqueous phase. The continuous phase, consisting of 2% fluorinated surfactant dissolved in FC-40 oil (FluoSurf_TM_, Emulseo) was injected with a syringe pump (TSE Systems) and a 1 mL glass syringe (BGB Analytik). Dispersed and continuous phases were connected to the chip with Tygon ND 100-80 tubing (Saint-Gobain, ID 0.5 mm). Aqueous solutions were co-flowed into a flow focusing junction along side with the dispersed phase. Droplet production was monitored using a 10X P-Apo air objective (NA 0.45) on a Nikon Ti-2E equipped with a SOLA SM II LED light source, a motorized stage and an Andor NEO 5.5 camera. Emulsions were recovered using a plastic syringe and incubated in the dark at 37°C, 250 rpm. The details of emulsions, their production and their analysis can be found in the supplementary information.

### High-throughput droplet scanning

For characterization, droplets were re-injected with a syringe pump (TSE Systems) into a dedicated chip treated similarly as the production chip. Spacer oil was injected with a 4-channel Elveflow OB1 controller. A white light laser source (SuperK Extreme, NKT Photonics) was used to excite fluorescence in the droplets at 491 and 565 nm through a Zeiss LD Achroplan 40x/0.60 Corr. Ph 2 microscopy objective at various rates (0.1-2kHz). Spacing oil flow through a flow-focusing junction allows droplets to be excited individually. The fluorescence emission signal was measured by PMTs (H10722-20, Hammmatsu) and recorded with a FPGA DAQ card (PCIE7841R, National Instruments) configured with LabVIEW. Laser beam positioning on the droplets was achieved using a high-speed camera (Mikrotron, Germany), an LED light source (M700L4, Thorlabs) and a custom-made microscope stage.

### Preparation of phage stock solution

Bacteriophage stocks of T7 phage were obtained by phage propagation on the host *E*.*coli* DSM 613 in liquid medium. Briefly, a overnight culture of *E*.*coli* was diluted 1:100 into 50 mL of NZCYM Medium (Carl Roth) and incubated at 37 °C, 250 rpm in a shaking incubator to OD 0.6-0.8. The culture was then infected with 10^8^ PFU of T7 phage and incubated for 3-4 h until full lysis. The solution was centrifuged 10 min, 7660 g to pellet bacteria debris and the supernatant was sterile-filtered using a 0.45 μm CFME syringe filter. Phage solution was supplemented with 0.1% BSA, aliquoted and stored at 4 °C. The titer of the phage solution was determined regularly by DLA to account for degradation. For each droplet experiment, the most current titer was used as reference.

### DLA

A double-layer agar plaque assay was used for a bulk determination of the phage titer. Liquefied top-agar (NCZYM, 0.75 % Agar) was mixed with 100 μl of bacterial culture in late-log or stationary phase and 100 μl of phages in varying dilutions, and the mixture was used to overlay bottom-agar (NZCYM, 1.5 % Agar). Plates were incubated at 37°C for at least 4 h until plaque formation was visible. The titer was determined as mean of all technical replicates. To determine titer development over time, DLA was performed with 2 different phage solution dilutions and plaques were counted at different timepoints. The titer for each timepoint was determined as mean of all replicates where plaques could be discerned. The absolute titer was determined as mean of all replicates from the latest timepoint where plaques could be distinguished for each of the dilutions.

### Bulk assay

Fluorescence emission intensity was measured in 384-well plates with a BMG CLARIOstar plate reader at 37°C and shaken at 700 rpm. Cell and phage solutions were diluted in PBS. All samples contained 1 μM YOYO-1 dye and T7 phages were added at a nominal 1.35 × 10^8^ PFU/mL.

## Data Availability

Source data supporting our findings is available in Zenodo at https://doi.org/10.5281/zenodo.11074305.

## Code Availability

Data analysis scripts are available in Zenodo at https://doi.org/10.5281/zenodo.11074305.

## Acknowledgments

This project has received funding by the European Commission through its Horizon 2020 research and innovation program under the Marie Skłodowska-Curie grant agreement No. 813786 (EVO-drops). We gratefully acknowledge funding by the Bavarian Ministry of Economic Affairs, Regional Development and Energy through an m4 award (project M4-2110-0004 “In vitro Synthese multivalenter Bakteriophagen zur Therapie von antibiotika-resistenten Infektionen”). We are thankful to Pr. Baret and David Van Assche for their assistance in the conception of our high-throughput droplet scanning setup.

## Author Contributions

L.G., S.v.S., F.K. conceptualised the project.

L.G. designed and performed experiments, wrote analysis scripts and wrote the original draft.

S.v.S. performed DLA and phage stock preparation.

F.K. wrote phage kinetics analysis scripts.

L.G., S.v.S., F.K., and F.C.S co-wrote the manuscript.

F.C.S supervised the work.

## Supplementary information

The online version contains supplementary material..

