## Supplementary Information for "Digital Phage Biology in Droplets"

L. Givelet, S. v. Schönberg, F. Katzmeier, F. C. Simmel

### 1 Baseline Model for Infection Kinetics

#### 1.1 Derivation

We consider a droplet containing $p$ phages and $b$ bacteria. To derive an expression for the probability over time that at least one phage in this droplet has infected a bacterium, we first consider the fate of a single encapsulated phage. The probability that this phage encounters a bacterium in a time interval $[t, t+dt]$ is proportional to $k\frac{b}{V}dt$, where $k$ is a constant and $\frac{b}{V}$ is the number density of the bacteria in the droplet. [NOTE: $\frac{p}{V}$, $\frac{b}{V}$ are the actual phage and bacterial densities in the droplet, while the bulk phage and bacterial densities are given by $c_p, c_b$].

We now denote $P_f(t)$ as the ('survival') probability of the phage not having infected a bacterium yet until time $t$, and $P_i(t)$ as the probability that the phage has already infected a bacterium. Over time, the survival probability of the phage drops, and the probability that an infection event has taken place rises correspondingly. This situation can be modeled with a time-continuous master equation in a two-state model for the single phage:

$$\frac{dP_f}{dt} = -k\frac{b}{V}P_f, \tag{1}$$

$$\frac{dP_i}{dt} = +k\frac{b}{V}P_f \tag{2}$$

At $t = 0$, we have the initial condition $P_f(t = 0) = 1$, and the solution of the first equation is thus given by

$$P_f(t) = e^{-k\frac{b}{V}t}. \tag{3}$$

We now consider all $p$ phages operating independently. The probability that none of the $p$ phages has infected a bacterium until time $t$ is simply given by $(P_f(t))^p$. For modeling our droplet experiments, we are interested in the occurrence of at least one infection, which is the complementary case. The probability $P_I$ of having at least one infection in a droplet at time $t$ can be thus be stated as

$$P_I = 1 - (P_f(t))^p = 1 - e^{-k\frac{b}{V}pt}. \tag{4}$$

We can now combine this result with the probabilities $P_b$ and $P_p$ of encapsulating exactly $b$ bacteria and $p$ phages, which follow a Poisson distribution with expected values $\lambda_b = (1-\alpha)c_bV$ and $\lambda_p = \alpha c_pV$ (cf. main text). Thus, the probability of finding a droplet with $p$ phages and $b$ bacteria, which has undergone a lysis event, is given by $P_p \cdot P_b \cdot P_I$. Summing over all possible droplet occupancies of phages and bacteria yields the probability $P_L(t)$ of finding a droplet where a phage has already infected a bacterium:

$$P_L = \sum_{b=0}^{\infty}\sum_{p=0}^{\infty} P_p \cdot P_b \cdot P_I \tag{5}$$

$$= \sum_{b=0}^{\infty}\sum_{p=0}^{\infty} \frac{\lambda_p^p}{p!}e^{-\lambda_p} \cdot \frac{\lambda_b^b}{b!}e^{-\lambda_b}\left(1 - e^{-k\frac{b}{V}pt}\right) \tag{6}$$

The above expression can be simplified by using $\sum_{p=0}^{\infty}\sum_{b=0}^{\infty} P_pP_b = 1$ and subsequently factoring out the $P_b$ terms from the inner sum:

$$P_L = 1 - \sum_{b=0}^{\infty} \frac{\lambda_b^b}{b!}e^{-\lambda_b} \sum_{p=0}^{\infty} \frac{\lambda_p^p}{p!}e^{-\lambda_p}e^{-k\frac{b}{V}pt} \tag{7}$$

When we substitute $s := e^{-k\frac{b}{V}t}$, we can identify the inner sum as the definition of the probability-generating function $M_P(s)$ of a Poisson distribution. Thus, we can replace the inner sum by the expression $M_P(s) = \exp(\lambda_p(s-1))$ and arrive at:

$$P_L = 1 - \sum_{b=0}^{\infty} \frac{(\lambda_b)^b}{b!}e^{-\lambda_b}e^{-\lambda_p(1-e^{-k\frac{b}{V}t})}. \tag{8}$$

As a side note, evaluating this function can cause numerical problems due to the potentially very large values of $b!$ and $(\lambda_b)^b$. An effective way to evaluate this sum is given by $P_L = 1 - \sum_{b=0}^{\infty} e^{Q_b}$, with $Q_b$ computed separately for each summand. $Q_b$ is given by

$$Q_b = b\ln\lambda_b - \lambda_b - \ln(\Gamma(b+1)) - \lambda_p(1 - e^{-k\frac{b}{V}t}) \tag{9}$$

where $\Gamma$ is the Gamma function, which should be evaluated numerically as the log Gamma function.

### 1.2 Approximation

We now proceed to find an approximate expression for the sum given in Eq. 8. According to the law of the unconscious statistician, the remaining sum can be interpreted as an expression for computing the expected value of a function $f(B)$, where $B$ represents the random number of encapsulated bacteria:

$$P_L = 1 - \mathbf{E}\left[e^{-\lambda_p\left(1-e^{-k\frac{B}{V}t}\right)}\right] =: 1 - \mathbf{E}\left[f(B)\right]. \tag{10}$$

For large expected values $\lambda_b$ of encapsulated bacteria, where $\lambda_b > 10$, the Poisson distribution can be approximated with a normal distribution with expected value $\lambda_b$ and standard deviation $\sigma = \sqrt{\lambda_b}$. Therefore, $P_L$ can be approximated as

$$P_L \approx 1 - \int_{-\infty}^{\infty} \frac{1}{\sqrt{2\pi\lambda_b}} \exp\left(-\frac{(b-\lambda_b)^2}{2\lambda_b}\right) f(b)db. \tag{11}$$

By substituting $x = (b-\lambda_b)/\sqrt{\lambda_b}$, we obtain the standard normal distribution and arrive at

$$P_L = 1 - \int_{-\infty}^{\infty} \frac{1}{\sqrt{2\pi}} \exp\left(-\frac{x^2}{2}\right) f(\sqrt{\lambda_b}x + \lambda_b)dx. \tag{12}$$

If we replace $f(\sqrt{\lambda_b}x + \lambda_b)$ with a Taylor series expansion around $x = 0$, we arrive at

$$P_L = 1 - \sum_{n=0}^{\infty} \int_{-\infty}^{\infty} \frac{1}{\sqrt{2\pi}} \exp\left(-\frac{x^2}{2}\right) f^{(n)}(\lambda_b) \frac{\left(\sqrt{\lambda_b}x\right)^n}{n!} dx. \tag{13}$$

In the above series, the odd terms in $x$ vanish since these integrals are overall odd, leaving us with

$$P_L = 1 - f(\lambda_b) - f''(\lambda_b)\frac{\lambda_b}{2} - \dots. \tag{14}$$

We now propose that a reasonable approximation for $P_L$ for large $\lambda_b$ is given by:

$$P_L \approx 1 - f(\lambda_b) = 1 - e^{-\lambda_p\left(1-e^{-k\frac{\lambda_b}{V}t}\right)} \tag{15}$$

To confirm this, we compute $f''(b)$ and compare its magnitude to that of $f(\lambda_b)$. The function and its derivatives are given by

$$f(b) = e^{-\lambda_p\left(1-e^{-\frac{kt}{V}b}\right)}, \tag{16}$$

$$f'(b) = e^{-\lambda_p\left(1-e^{-\frac{kt}{V}b}\right)} e^{-\frac{kt}{V}b} \lambda_p \frac{kt}{V}, \tag{17}$$

$$f''(b) = e^{-\lambda_p\left(1-e^{-\frac{kt}{V}b}\right)} \left(\frac{kt}{V}\right)^2 e^{-\frac{kt}{V}b} \left(\lambda_p e^{-\frac{kt}{V}b} + 1\right) \lambda_p. \tag{18}$$

The magnitude of both terms can be compared by computing

$$\delta = \frac{f''(\lambda_b)\frac{\lambda_b}{2}}{f(\lambda_b)} = \frac{\lambda_b}{2}\left(\frac{kt}{V}\right)^2 e^{-\frac{kt}{V}\lambda_b}(\lambda_p e^{-\frac{kt}{V}\lambda_b}+1)\lambda_p. \tag{19}$$

This expression is 0 at $t = 0$ and approaches 0 as $t \to \infty$. Further, the term $\left(\frac{kt}{V}\right)^2 e^{-\frac{kt}{V}\lambda_b}$ has a single maximum at $\frac{kt}{V} = \frac{2}{\lambda_b}$ with the value $\frac{4}{e^2\lambda_b^2}$. Thus, we can conclude that

$$\delta = \frac{f''(\lambda_b)\frac{\lambda_b}{2}}{f(\lambda_b)} < \frac{2}{e^2}\frac{\lambda_p}{\lambda_b}(\lambda_p e^{-\frac{kt}{V}\lambda_b}+1), \tag{20}$$

which provides an estimate of the accuracy of our approximation and validates the initial assertion that the approximation becomes accurate for large $\lambda_b$. For conditions similar to our experimental conditions ($\lambda_p = 2$ and $\lambda_b = 10$), we find that $\delta < 0.054$ for large enough $t$. As a further validation, we plotted the full $P_L(t)$ from equation 8 and our approximation from equation 15 for several values of $\lambda_p$ and $\lambda_b$ in Figure S1. We find that the approximation is fairly accurate for most values of $\lambda_b \geq 10$ and $\lambda_p \leq 5$.

Equation 15 can be rewritten using $\lambda_p = \alpha c_p V$ and $\lambda_b = (1-\alpha)c_b V$, resulting in

$$P_L \approx 1 - e^{-\alpha c_p V\left(1-e^{-(1-\alpha)kc_b t}\right)} =: 1 - e^{-\alpha c_p V A(t)}, \tag{21}$$

which corresponds to the form given in the main text.

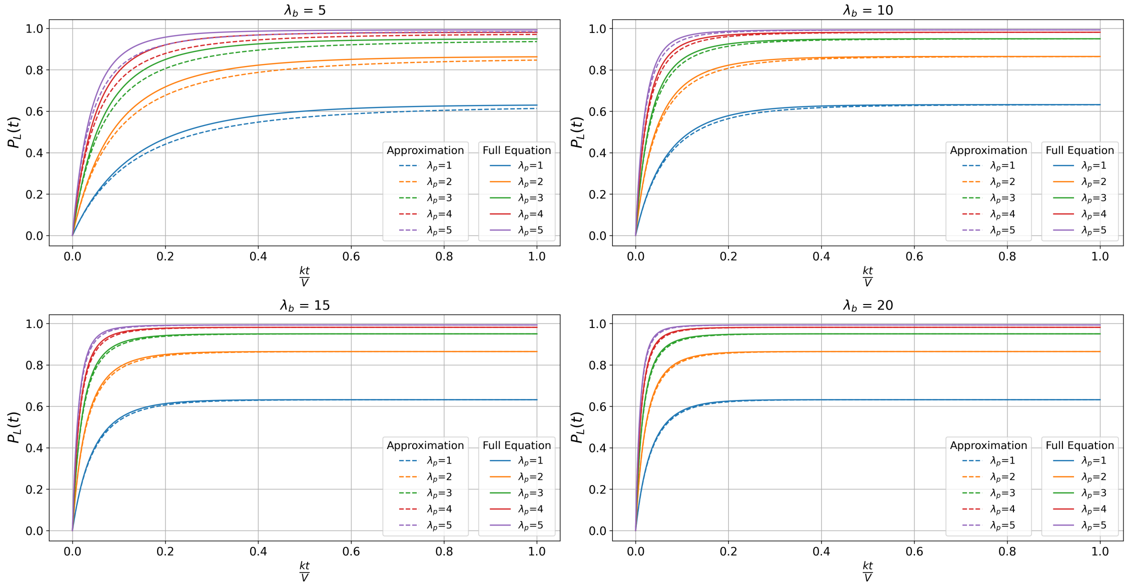


**Fig. S1** Plot of the approximated and full equation of the probability $P_L(t)$ of finding a droplet where at least one phage has infected a bacterium over time.

### 2 Emulsion production

To produce emulsions with different mixing ratios, alternating pressures were applied to the aqueous phases in cycles of 30 seconds while keeping the total pressure $p_tot$ applied to the dispersed phase constant. For the production of emulsions with a continuous mixing ratio variation, two opposing linear pressure ramps with constant total pressure were applied. The red fluorescence channel was used to determine the mixing ratio $\alpha$ during production. We assumed $\alpha = 0.5$, when the pressures applied to the aqueous phases are equal. To produce an emulsion with a bimodal size distribution, the total pressure applied to the aqueous phases was alternated in cycles of 30 seconds. The bright-field channel was used to measure the droplet diameters through the auto-segmentation feature of the NIS-Elements software. For emulsions exhibiting a bimodal size distribution, a microfluidic device with an observation section located downstream of the flow focusing junction with a channel height considerably higher than droplet diameter (h > $100\,\mu\mathrm{m}$) was used. This ensured the droplets were spherical, allowing for accurate diameter measurements. The flow rate of the continuous phase was maintained constant during each droplet production session, but this rate was varied from session to session in the range from 1100 to 2100 µL/h.

### 3 Droplet analysis

To analyse droplets, a dedicated custom-made python script was used. Briefly, each droplet is characterized by a red and green fluorescence signal height and width. To exclude any outlier, droplets are filtered based on their size and fluorescence signal, and time of acquisition through arbitrary threshold values entered by the user.

For emulsions containing different $\alpha$, droplets were classified by red PMT fluorescence signal height. For emulsions containing several size modes, droplet were classified in function of the red fluorescence signal width. The $\alpha$ values and droplet diameters determined through microscope image analysis were attributed to the dominant modes of red signal height and width, respectively. Digitization was carried out using the machine learning Python package scikit-learn [1]. A Gaussian Mixture Model (GMM) was applied to the logarithm-transformed green fluorescence signal values, considering 2 to 5 components (3 to 5 for continuous $\alpha$ screening). The silhouette score [2] was computed for each component number and the model configuration with the highest score was selected. The fraction of positive droplets was determined by summing the weights of the highest modes predicted by the GMM. For emulsion comprising a continuous variation of $\alpha$ values, GMM analysis was applied to each bin. Bacteria droplet occupancy $\lambda_b$ were calculated using droplet volume $V$ and bacterial density $c_b$. The latter was computed from OD bulk measurements, with the standard correlation used for *E. coli* i.e. 1 unit of $OD_{600}$= $8 \times 10^8\,\mathrm{cells/mL}$.

| OD | DLA Titer (PFU/mL) | $\alpha$ | Droplet diameter (µm) | $\lambda_b$ (cells/droplet) | Digitized droplets | Silhouette Score | $P_L$ |
|---|---|---|---|---|---|---|---|
| 2.5 | $2.50 \times 10^7 \pm 9.74 \times 10^6$ | 0.5 | 37.622 ± 0.731 | 27.882 | $6.69 \times 10^4$ | 0.733 | 0.299 |
| 2.5 | $2.50 \times 10^8 \pm 9.74 \times 10^7$ | 0.5 | 37.074 ± 0.755 | 26.681 | $1.01 \times 10^5$ | 0.643 | 0.868 |

**Table 1** Details of emulsions used in Figure 2. Droplets were scanned after approximately 90 min of incubation. Silhouette Score is evaluated on the droplet clusters predicted by GMM.

| OD | DLA Titer (PFU/mL) | $\alpha$ | Droplet diameter (µm) | $\lambda_b$ (cells/droplet) | Digitized droplets | Silhouette Score | $P_L$ |
|---|---|---|---|---|---|---|---|
| 3.2 | $9.25 \times 10^7 \pm 5.30 \times 10^6$ | 0.260 | 47.61 ± 0.970 | 107.045 | $1.09 \times 10^4$ | 0.616 | 0.782 |
| 3.2 | $9.25 \times 10^7 \pm 5.30 \times 10^6$ | 0.353 | 47.61 ± 0.970 | 93.592 | $9.30 \times 10^3$ | 0.609 | 0.728 |
| 3.2 | $9.25 \times 10^7 \pm 5.30 \times 10^6$ | 0.608 | 47.61 ± 0.970 | 56.705 | $2.01 \times 10^4$ | 0.612 | 0.580 |
| 3.2 | $9.25 \times 10^7 \pm 5.30 \times 10^6$ | 0.682 | 47.61 ± 0.970 | 46.000 | $7.59 \times 10^3$ | 0.605 | 0.517 |
| 3.2 | $9.25 \times 10^7 \pm 5.30 \times 10^6$ | 0.764 | 47.61 ± 0.970 | 34.139 | $6.86 \times 10^3$ | 0.609 | 0.318 |

**Table 2** Details of emulsions used in Figure 3. Droplets were scanned after approximately 240 min of incubation. Silhouette Score is evaluated on the droplet clusters predicted by GMM.

| OD | DLA Titer (PFU/mL) | $\alpha$ | Droplet diameter (µm) | $\lambda_b$ (cells/droplet) | Digitized droplets | Silhouette Score | $P_L$ |
|---|---|---|---|---|---|---|---|
| 4.65 | $1.18 \times 10^7 \pm 1.77 \times 10^6$ | Min: 0.368<br>Max: 0.936 | 37.03 ± 0.96 | Min:6.330<br>Max: 62.506 | $7.20 \times 10^5$ | Min:0.520<br>Max: 0.725 | Min: 0.559<br>Max:0.938 |

**Table 3** Details of emulsions used in Figure 4. Droplets were scanned after approximately 140 min of incubation. Silhouette Score is evaluated on the droplet clusters predicted by GMM.

| OD | DLA Titer (PFU/mL) | $\alpha$ | Droplet diameter (µm) | $\lambda_b$ (cells/droplet) | Digitized droplets | Silhouette Score | $P_L$ |
|---|---|---|---|---|---|---|---|
| 4.40 | $9.55 \times 10^7 \pm 2.07 \times 10^6$ | 0.5 | $34.03 \pm 0.96$ | 36.316 | $t_1 : 1.32 \times 10^5$<br>$t_2 : 3.51 \times 10^4$<br>$t_3 : 1.36 \times 10^5$<br>$t_4 : 4.79 \times 10^4$<br>$t_5 : 2.73 \times 10^4$<br>$t_6 : 2.41 \times 10^4$<br>$t_7 : 6.10 \times 10^3$<br>$t_8 : 1.91 \times 10^5$ | $t_1 : 0.810$<br>$t_2 : 0.757$<br>$t_3 : 0.768$<br>$t_4 : 0.721$<br>$t_5 : 0.725$<br>$t_6 : 0.665$<br>$t_7 : 0.720$<br>$t_8 : 0.609$ | $t_1 : 0.119$<br>$t_2 : 0.214$<br>$t_3 : 0.324$<br>$t_4 : 0.340$<br>$t_5 : 0.391$<br>$t_6 : 0.453$<br>$t_7 : 0.460$<br>$t_8 : 0.568$ |
| 4.40 | $9.55 \times 10^7 \pm 2.07 \times 10^6$ | 0.5 | $43.95 \pm 1.03$ | 78.233 | $t_1 : 3.16 \times 10^5$<br>$t_2 : 1.22 \times 10^5$<br>$t_3 : 4.90 \times 10^5$<br>$t_4 : 4.45 \times 10^4$<br>$t_5 : 6.54 \times 10^4$<br>$t_6 : 8.10 \times 10^4$<br>$t_7 : 7.10 \times 10^4$<br>$t_8 : 4.35 \times 10^5$ | $t_1 : 0.745$<br>$t_2 : 0.674$<br>$t_3 : 0.672$<br>$t_4 : 0.620$<br>$t_5 : 0.611$<br>$t_6 : 0.559$<br>$t_7 : 0.602$<br>$t_8 : 0.550$ | $t_1 : 0.201$<br>$t_2 : 0.320$<br>$t_3 : 0.470$<br>$t_4 : 0.526$<br>$t_5 : 0.602$<br>$t_6 : 0.727$<br>$t_7 : 0.733$<br>$t_8 : 0.807$ |

**Table 4** Details of emulsions used in Figure 5. Exposure time for each emulsion are indicated. $t_1$ = 81 min, $t_2$ = 137 min, $t_3$ = 191 min, $t_4$ = 263 min, $t_5$ = 315 min, $t_6$ = 406 min, $t_7$ = 499 min, $t_8$ = 1100 min.Silhouette Score is evaluated on the droplet clusters predicted by GMM.

### Mixing ratio mapping

In contrast to discrete mixing ratio screening, the continuous variation of mixing fraction $\alpha$ produced a smooth distribution of red fluorescence values. Through microscopy analysis we determined the smallest and highest observed mixing ratio (respectively $\alpha_{min}$ and $\alpha_{max}$). We linearly mapped bins to $\alpha$ values to by using $\alpha_{min}$ and $\alpha_{max}$ as reference points. The $\alpha_{min}$ and $\alpha_{max}$ determined through microscopy were 0.368 and 0. 936, respectively, based on the analysis of 1101 intensity profiles (Figure S5).

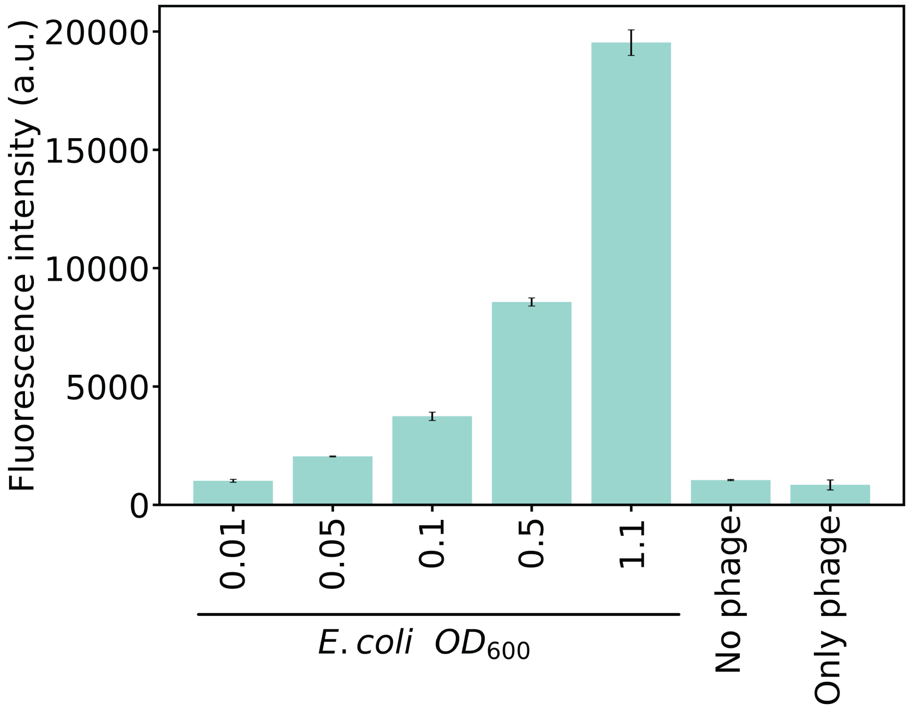


**Fig. S2** Bacterial lysis and DNA intercalation. Bars represent YOYO-1 fluorescence emission intensity measured 110 min after phage addition. Errors bars represent standard deviation over technical replicates, n=3. Cells alone and phage alone display similar fluorescence levels, through unspecific cell lysis and phage DNA labelling, respectively.

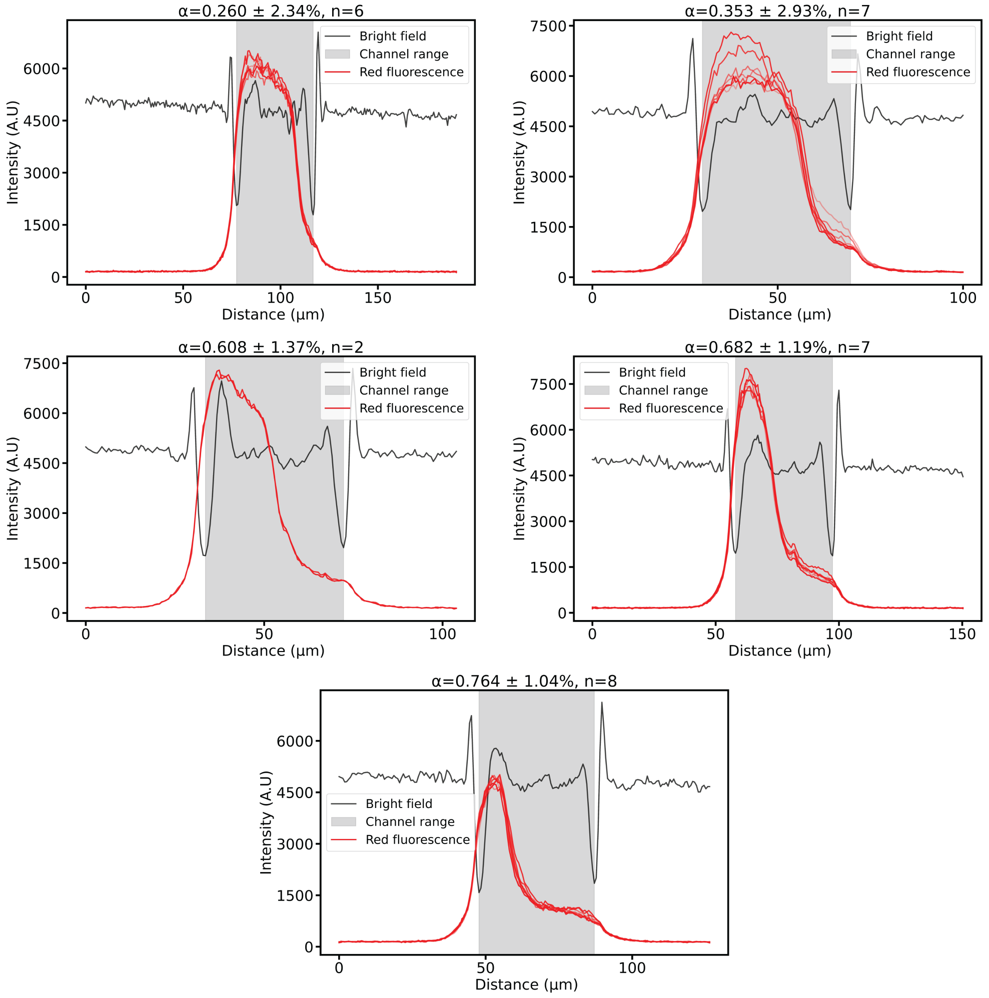


**Fig. S3** Determination of $\alpha$ by image analysis. Intensity profiles are sampled across the channel containing the two laminar streams of aqueous solutions. The intensity profile of bright-field images is used to determine the microfluidic channel range. The fluorescence value at the left bound of the channel serves as the threshold to identify significant fluorescence signal within the channel. The extent of the fluorescence signal within the channel is divided by the total channel length to determine the proportion of the fluorescent solution in the total aqueous phase. Since the reference dye is present in the solution containing bacteria, the $\alpha$ value equals one minus the fluorescent proportion in the aqueous phase.

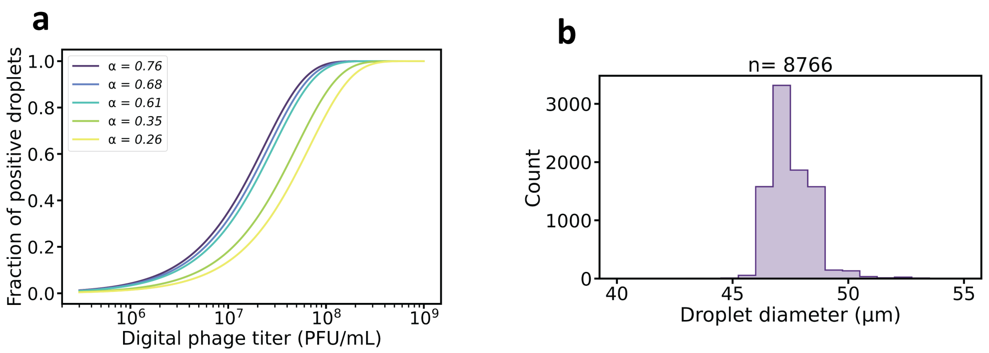


**Fig. S4** Emulsion characterisation through microscopy of the emulsion used for discrete $\alpha$ screening. **(a)** Relation between positive droplet fraction and digital titre for droplet with the obtained $\alpha$ values. Each mixing ratio mode exhibits a substantial difference in its optimal dynamic range.**(b)** Distribution of droplet sizes determined through image analysis, exhibiting an average diameter of 47.61 ± 0.97 µm. During production, mode of larger droplets caused by coalescence in the chip was excluded from analysis, both in droplet acquisition and image analysis.

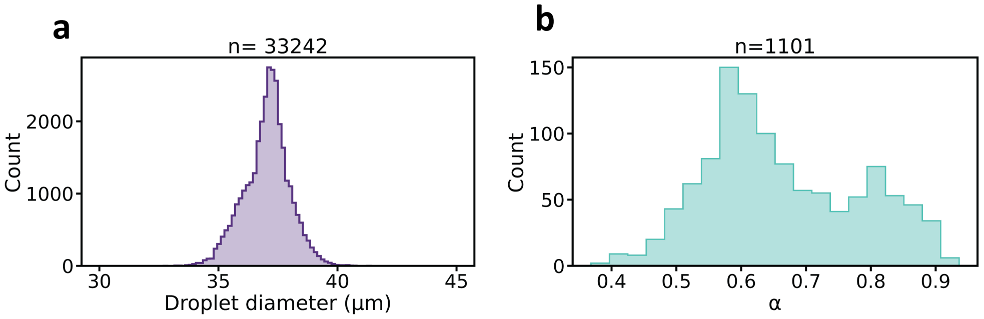


**Fig. S5** Emulsion characterisation through microscopy of the emulsion used for continuous $\alpha$ screening. **(a)** Distribution of droplet sizes determined through image analysis, exhibiting an average diameter of 37.03 ± 0.96 µm. **(b)** $\alpha$ values determined through image analysis. Mixing ratios were determined using a method closely resembling the one described in Figure S3, with the minor distinction that here, the proportion of the fluorescence signal was measured at the focusing junction.

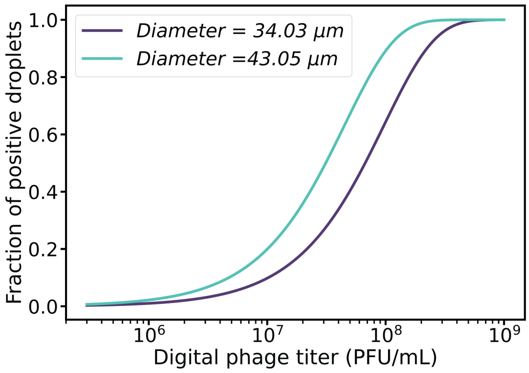


**Fig. S6** Relation between positive droplet fraction and digital titre for two different droplet size. Droplet sizes exhibit a substantial difference in their optimal dynamic range.

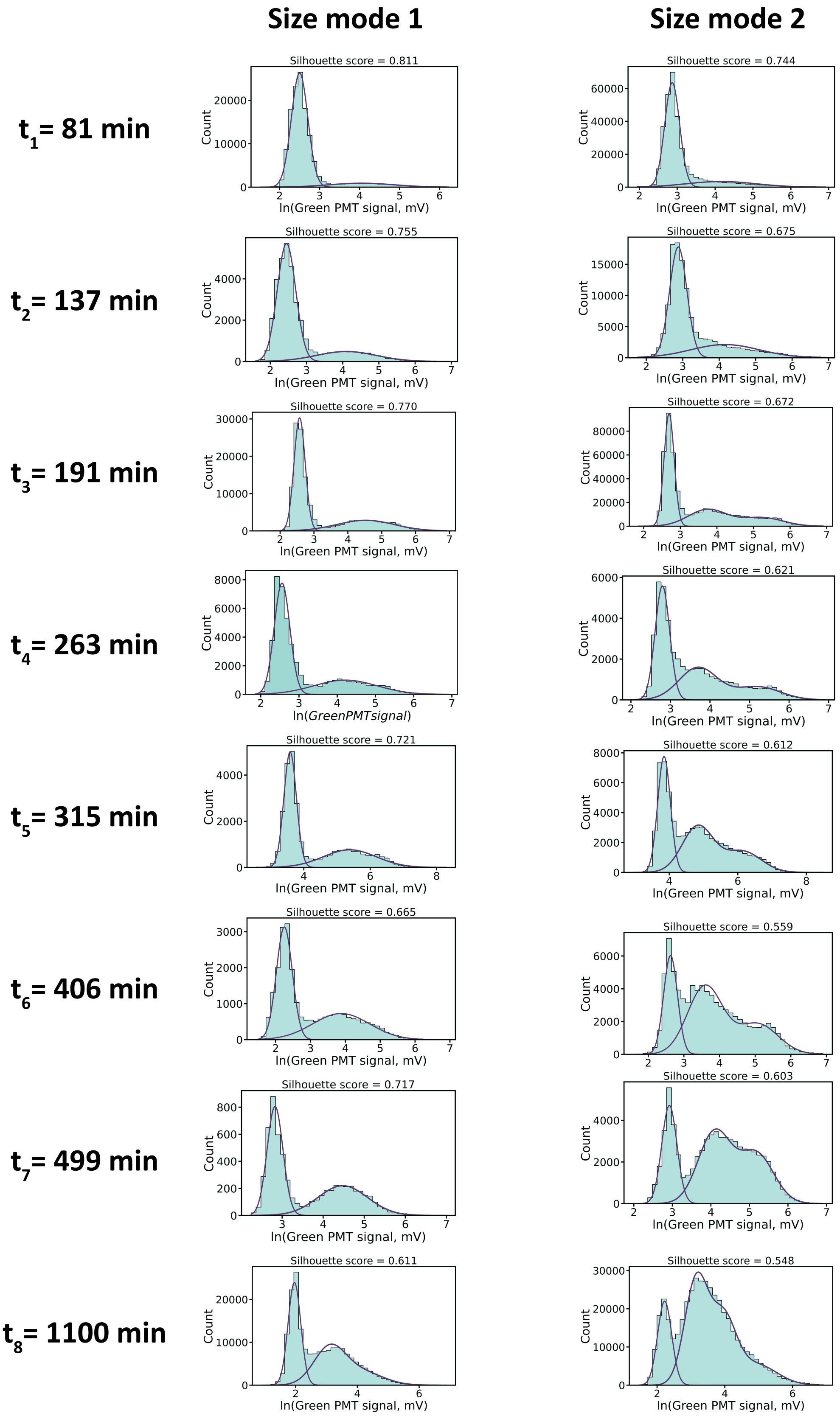


**Fig. S7** Droplet digitization for different size and exposure times. Blue bars represent the logarithm-transformed green fluorescence signal values and the purple lines positives and negative droplets modes predicted by GMM with the associated silhouette score indicated on top of each plot.
